# The neural organization of visual information in the auditory cortex of the congenitally deaf

**DOI:** 10.1101/2023.04.17.537188

**Authors:** Zohar Tal, Joana Sayal, Fang Fang, Yanchao Bi, Jorge Almeida, Alessio Fracasso

**Affiliations:** Proaction Laboratory, Faculty of Psychology and Educational Sciences, University of Coimbra, Portugal; Center for Research in Neuropsychology and Cognitive and Behavioral Intervention (CINEICC), Faculty of Psychology and Educational Sciences, University of Coimbra, Portugal; School of Psychological and Cognitive Sciences and Beijing Key Laboratory of Behavior and Mental Health, Peking University, Beijing, China; IDG/McGovern Institute for Brain Research, Peking University, 100087 Beijing, China; Peking-Tsinghua Center for Life Sciences, Peking University, 100087 Beijing, China; State Key Laboratory of Cognitive Neuroscience and Learning and IDG/McGovern Institute for Brain Research, Beijing Normal University, Beijing, China; Beijing Key Laboratory of Brain Imaging and Connectomics, Beijing Normal University, Beijing, China; Chinese Institute for Brain Research, Beijing, China; School of Neuroscience and Psychology, University of Glasgow, Scotland

**Keywords:** Neuroplasticity, Functional Magnetic Resonance Imaging, Population Receptive Field Analysis, Congenital Deafness, Topographic Organization

## Abstract

Neuroplasticity is the ability of the human brain to reorganize and modify its activity throughout life. In congenital deafness, sensory-deprived cortex can be recruited to represent sensory information belonging to other modalities, a process known as cross-modal plasticity. Previous studies have indicated that the auditory cortex of congenitally deaf, but not of hearing individuals, is recruited during visual tasks. However, it is not clear to what extent, and how, these cross-modal responses in the deprived auditory cortex represent low-level visual spatial information or map the visual field. Here, we addressed this question directly in an fMRI case-study, aiming to map retinotopic features in the auditory cortex. Two congenitally deaf and one hearing participant went through a conventional retinotopy fMRI experiment with visual stimuli designed to map the visual system. Using population receptive field (pRF) modelling, we revealed retinotopic-related responses in the auditory cortex of the deaf, but not in the hearing. These responses, that were mostly lateralized to the right hemisphere, represented the contralateral visual field, and were characterized by large receptive fields, centred to near foveal areas. Interestingly, we found that these responses to visual stimuli predominantly reflected negative BOLD signals in the auditory cortex of the deaf, suggesting that visual information might be represented through cross-modal deactivation signals.

## Introduction

The human brain can partially reorganize and modify its activity throughout life (Frasnelli et al., 2011). Psychophysical and imaging methods have provided evidence in relation to the development of human visual system (Kiorpes, 2016) reaching the conclusion that the visual pathways up through primary visual cortex are remarkably mature already in infants (Kiorpes, 2016). However, evidence also suggests that behaviourally relevant phenomena can emerge at a later stage in human development (Carrasco et al., 2022), which is compatible with the existence of environmental factors shaping the developing visual system. Crucially, lesion and sensory deprivation studies represent an important line of study, indicating that human early visual cortex has the ability to substantially reshape itself following sensory deprivation, showing examples of neuroplasticity (Collignon et al., 2009; Gougoux et al., 2004; Van Boven et al., 2000).

One major example of neuroplasticity comes from the study of sensory deprivation in case of congenital blindness or deafness. Under these circumstances, the sensory-deprived cortex can be recruited to process information belonging to other, intact modalities, a process known as cross-modal plasticity. Cross-modal neuroplasticity has been extensively studied in animal and human blindness (Collignon et al., 2009; Gougoux et al., 2004; Van Boven et al., 2000) revealing the involvement of the deprived visual cortex in non-visual tasks including auditory, tactile, and language processing (Amedi et al., 2003; Bedny et al., 2011; Collignon et al., 2009; Ptito et al., 2012; Renier et al., 2010; Sadato et al., 1996). Similarly, a substantial body of research has documented the recruitment of auditory-deprived areas during visual or somatosensory stimulation (Almeida et al., 2015; Bola et al., 2017; Bottari et al., 2014; Karns et al., 2012; Scurry et al., 2020; for review see Bell et al., 2019). Which principles govern the organization of sensory information in cross-modal plasticity? Do cross-modal responses in deprived cortical areas represent low-level information in a similar way to the intact sensory areas? Here, we explored the representation of low-level spatial visual information in the auditory cortex of deaf individuals.

Functional neuroimaging studies show that the auditory cortex (AC) of congenitally deaf individuals reorganizes to process visual information, such as visual motion (Retter et al., 2018), rhythm/frequency discrimination (Bola et al., 2017) and position in the visual field (Almeida et al., 2015). Cross-modal plasticity was also documented in electrophysiological studies with animal models of deafness, including cats (Land et al., 2016; Lomber et al., 2010, 2011; Meredith et al., 2011; Rebillard et al., 1977); ferrets (Meredith et al., 2012; Meredith & Allman, 2012) and mice (Hunt et al., 2006). Congenital or early auditory deprivation was also associated with behavioural changes, such as an improved performance in various sensorial tasks of the unaffected modalities. For example, deaf individuals demonstrate enhanced visual abilities in motion detection (Almeida et al., 2018; Hauthal et al., 2013) and visual localization tasks (Codina et al., 2011; Seymour et al., 2017), especially when the stimuli are presented in the peripheral visual field (Bavelier et al., 2000; Neville & Lawson, 1987). Congenitally deaf cats show superior performance in visual localization in peripheral visual fields and lower visual motion detection thresholds compared to hearing cats (Lomber et al., 2010, 2011). These attributes can easily be understood as compensatory mechanisms for the auditory loss because these functions in hearing individuals work in tandem – i.e., auditory and visual input - for the perception of daily life danger and spatial orienting.

Animal studies can elucidate some of the mechanisms underlaying these processes, by localizing individual visual functions to portions of the reorganized AC and establishing a causal relationship between the activated areas and visual performance (Lomber et al., 2010; Meredith et al., 2011; Meredith & Lomber, 2011). For example, reversible deactivation of the dorsal zone (DZ) in cats reduced their enhanced motion detection abilities while deactivation of the posterior auditory field (PAF) abolished superior visual peripheral localization (Lomber et al., 2010). These findings suggest that cross-modal processes might underlie the superior performance of deaf individuals in different tasks related to the remaining senses. Moreover, single-unit recordings in deaf (or partially deaf) animals have shown an increase in the proportion of neurons in the auditory cortex that respond to visual and tactile stimuli (Hunt et al., 2006; Meredith et al., 2011, 2012; Meredith & Lomber, 2011). Detailed characterization of the visual-evoked responses of these neurons revealed large-diameter receptive fields, which were distributed across the contralateral visual hemifield, extending also to the ipsilateral hemifield (Meredith et al., 2011; Meredith & Lomber, 2011).

Does the reorganized auditory cortex of humans represent spatial features of visual stimuli in a similar way? In a study by Almeida et al. (2015), a group of deaf and hearing individuals was presented with a set of stimuli typically used to map visual field location. Using multi-voxel pattern analysis, the authors found that the location of the visual stimuli could be decoded from the activity pattern of the primary auditory cortex of the deaf, but not the hearing group. These results clearly show that the activity pattern of the primary auditory cortex of deaf individuals contains spatial visual information. However, a detailed examination of visual receptive fields’ properties in the deprived auditory cortex has not been performed so far.

Following Almeida et al. (2015), here, we addressed this question directly, by characterizing visual responses in the AC of deaf individuals and exploring their spatial organization using population receptive field (pRF) modelling (Dumoulin & Wandell, 2008). In pRF analysis, the collective response and tuning properties of the neuronal population are modelled at each recording site (voxel), providing estimation of receptive fields’ location and size (tuning width). pRF modelling, which was first introduced for retinotopic mapping (Dumoulin & Wandell, 2008) has been extensively used to map other sensory cortices and higher-order areas (Fracasso, Petridou, et al., 2016; Harvey et al., 2013; Scurry et al., 2020; Thomas et al., 2015). Here, we use pRF modelling to provide a descriptive account of the properties characterizing cross-modal visual receptive fields in humans.

## Methods

### Participants

One hearing individual (female, 19 years old) and two congenitally deaf individuals (2 females, 17 and 21 years old) participated in the experiment; all were naive to the purpose of the experiment. Participants had normal or corrected-to-normal vision, no history of neurological disorder, and gave written informed consent in accordance with the guidelines of the institutional review board of Beijing Normal University Imaging Center for Brain Research. Both deaf participants were proficient in Chinese sign language and had hearing loss above 90 dB binaurally (frequencies tested ranged from 125 to 8,000 Hz). One of them never used hearing aids, and the other used a hearing aid on her left ear every day for 10 years since she was 5 years old, which allowed her to perceive sound, but not discriminate between different categories (e.g., voices) or localize its source in space. The causes of deafness in both participants were pregnancy-related complications. The hearing participant reported no hearing impairment or knowledge of Chinese sign language.

### Stimuli and procedure

The stimuli used in the experiment were generated in MATLAB (MathWorks) using the PsychToolbox (Brainard, 1997; Kleiner et al., 2007; Pelli, 1997). Participants were presented with two types of visual stimuli that are typically used to obtain visual field maps: rotating wedges and expanding annuli. The stimulus field of view covered 6 degrees of visual angle (radius). Stimulation consistent in two separate stimuli. The first stimulus was a counterphase flickering (5 Hz) checkerboard wedge (30°) rotating along 12 equidistant positions along the 360°, eliciting different polar angle preferences (**Figure 1A**). The second type of stimuli were counterphase flickering (5 Hz) checkerboard annuli expanding in 12 different steps, starting from the central fovea up to the periphery of the visual field, eliciting eccentricity preferences (**Figure 1B**). Participants were asked to maintain fixation on a central point throughout the entire functional runs. Participants completed twelve runs: six in which the wedge stimuli rotated in clockwise order, starting from the top vertical plane, and six in which the annuli appeared with increasing radius. Each run (with a total of 84 volumes) started with 12 sec of blank (grey) screen, followed by six cycles of the full loop of stimuli (72 volumes) and ended with a blank screen for 12 sec.

**Figure 1:**
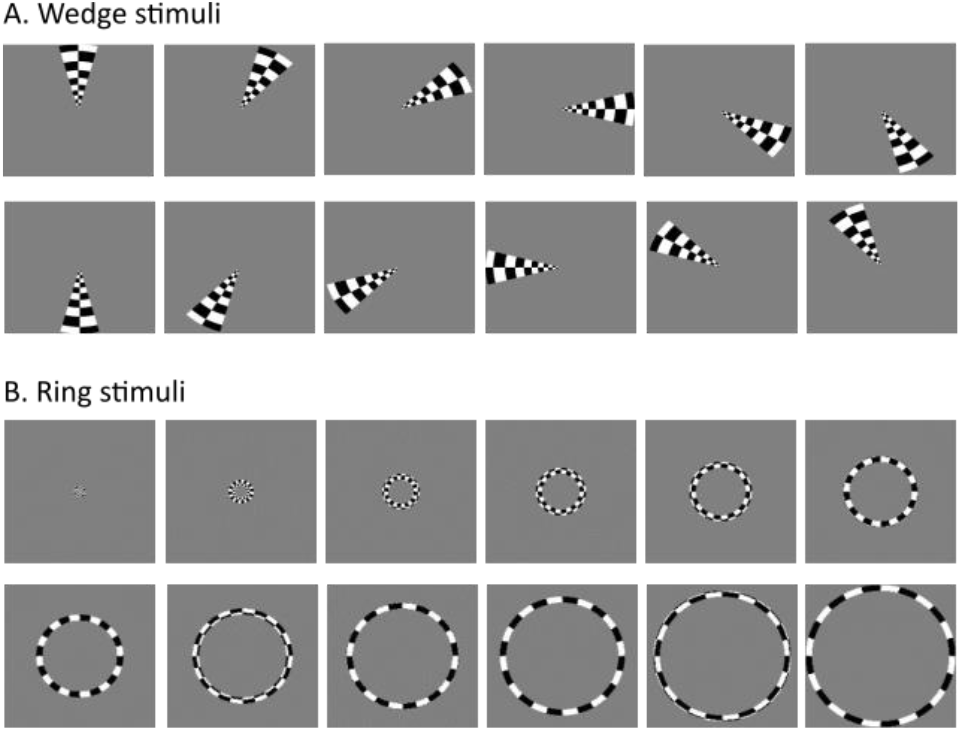
Experimental stimuli and procedure. **(A)** Polar angle stimuli. High contrast, flickering checkerboard wedges extending from the fovea to the visual periphery were presented along twelve polar angles (each wedge was presented for 1 TR= 2 sec). The wedge stimulus rotated clockwise around a central fixation point, eliciting each voxel’s preferred polar angle in the visual space. **(B)** Eccentricity stimuli. These flickering checkerboard stimuli elicited each voxel’s preferred eccentricity by presenting an expanding ring starting from the central fovea up to the periphery, in twelve different positions around a central fixation point (each ring was presented for 1 TR= 2 sec).

### Anatomical and Functional Imaging

magnetic resonance imaging (MRI) data was collected at the Beijing Normal University MRI center, on a 3 T Siemens Tim Trio scanner. A high-resolution 3-D structural data set was collected in the sagittal plane with a 3-D magnetization-prepared rapid-acquisition gradient echo sequence and the following parameters: repetition time (TR) = 2530 ms, echo time (TE) = 3.39 ms, flip angle = 7°, matrix size = 256×256, voxel size = 1×1×1.33 mm, 144 slices, acquisition time = 8.07 min. An echo-planar image sequence was used to collect functional data (TR = 2000 ms, TE = 30 ms, flip angle = 90°, matrix size = 64×64, voxel size = 3.125×3.125×4 mm, 33 slices, interslice distance = 4.6 mm, slice orientation = axial).

### fMRI pre-processing

Processing of the anatomical T1-weighted MR images was performed in Freesurfer image analysis suite (6.0 version, http://surfer.nmr.mgh.harvard.edu/) using the recon-all pipeline, that included white and grey matter segmentation, skull removal, and cortical reconstruction. Pre-processing of functional MRI data was performed in AFNI (Cox, 1996; Cox & Hyde, 1997) and included the following steps: slice acquisition time correction, head motion correction, and detrending and scaling. Following the initial pre-processing steps, the time-series of each stimulus type (wedge and ring) were averaged across repetitions for each participant and concatenated to create a single time-series for each participant, containing both polar angle and eccentricity stimuli. The averaged functional image was co-registered to the anatomical scans, using the function 3dAllineate (with local Pearson correlation as the cost function).

### Population receptive field analysis

Population receptive field (pRF) modelling (Dumoulin & Wandell, 2008) was conducted in R (R Core Team, 2014) following previous retinotopic mapping procedures (Dumoulin & Wandell, 2008; Fracasso, Petridou, et al., 2016). Briefly, a set of predicted pRF time-series were generated using a 2D-Gaussian model with three parameters-x, y (horizontal and vertical coordinates of the receptive field’s centre, based on a 28×28 grid of preferred visual locations) and σ (pRF size, ranging between 0.5 to 11 degrees of the visual field). The predicted time-series were calculated by convolving the stimulus sequence of a given pRF with a canonical hemodynamic response function (HRF, (Friston et al., 1998), parameters: 6-sec peak latency and 12-sec decay latency). For each voxel, the best fitting pRF was obtained by iteratively fitting all predicted pRFs time-series by means of a general linear model (GLM), storing the predicted pRF that yielded the highest variance explained (R^2^) with the constraint of a positive beta value (“positive model”). The extracted model parameters included x and y position, pRF size as well as the model slope (beta value), polar angle, eccentricity, and variance explained (R^2^). Variance explained values were used to threshold the data for visualization of the maps that were projected on the reconstructed cortical surface of each participant through AFNI/SUMA (Saad et al., 2004; Saad & Reynolds, 2012). In the visual areas, the threshold was set to 20% of variance explained. In the whole-brain or auditory areas analyses the threshold was set to 8% of variance explained. This value was set to be higher than the mean variance explained obtained from the white matter in all three participants. Mean white matter R^2^ values were: 0.06 ±0.09 for the hearing control (HC) participant; 0.08 ±0.13 for participant deaf 01 (D1); and 0.06 ±0.1 for participant deaf 02 (D2). It is important to note that the pRF method and its different implementations have been successfully used in the past to provide evidence about the plasticity and the stability of the visual system in case of congenital (Fracasso, Koenraads, et al., 2016) and acquired deficits (Levin et al., 2010).

Here we also included a model for the negative fluctuations of the blood-oxygen-level-dependent (BOLD) signal, where we removed the constraint of a positive beta value, which enabled to explore “negative pRFs” (“negative model”). Most of the research on cross-modal processing, and particularly in the context of congenital sensory deprivation, focused on positive BOLD responses, searching for sensory-specific areas showing increased activity in response to stimuli from other modalities. However, cross-modal stimuli could also induce negative BOLD responses (NBR, also referred as deactivation) in sensory-specific cortical areas (Hairston et al., 2008; Jorge et al., 2018; Laurienti et al., 2002; Merabet et al., 2007; Sadato et al., 1996; Tal et al., 2016; Weisser et al., 2005) or in other, non-sensory specific areas such as the default mode network (DMN) (Anticevic et al., 2012; Mayer et al., 2009). Recently, it has been shown that NBR represent spatial visual information, and that the time-series of this deactivations could be modelled using pRF methods (Szinte & Knapen, 2020). Here, we applied a similar approach to study the visual organization of cross-modal processing in the auditory cortex of deaf.

In a traditional pRF modelling, the best-fitting predictor is determined based on the highest variance explained (R^2^) and positive beta parameter. Here, in the model that included negative pRF, we removed the constraint on the positive beta value, and extracted the model parameters for the pRF that yielded the highest variance explained (R^2^). In this way the best fitting time series could be characterized by either positive or negative beta values.

### Region of interest definition

To extract the models’ parameters and compare between the positive and negative models, we defined individual regions of interest (ROI) in visual and auditory areas, as well as a control ROI in the white matter. White matter ROI was defined based on the individual segmentation pipeline of Freesurfer (6.0 version, http://surfer.nmr.mgh.harvard.edu/). Visual and auditory ROIs were defined based on the Human Connectome Project (HCP) multimodal parcellation (MMP) atlas (Glasser et al., 2016) available in AFNI/SUMA. The cortical atlas was projected on the standard cortical surface of each individual participant, and then resampled to volumetric space. Model parameters were extracted for all the voxels in a chosen ROI. For statistical analysis, individual ROIs were grouped into three main regions: early visual areas, early auditory areas, and higher auditory areas (following the grouping criteria in Glasser et al. (2016). Early visual ROI included V1, V2 and V3 cortical areas; Early auditory ROI included A1, medial belt, lateral belt, para belt and retro-insular cortical areas; Higher auditory ROI included A4, A5, STSdp, STSda, STSvp, STSva, STGa, and TA2.

### Bootstrapping analysis

To compare the goodness of fit of the positive and negative pRF models we performed a bootstrapping analysis of the variance explained values in the early visual cortex and the white matter ROIs. To test for potential differences between the models at a wide range of variance explained thresholds, bootstrapping was performed using quantile thresholding for each participant. For that, within each ROI, voxels were selected based on variance explained values, thresholded in 5 quantile steps (ranging between 0.3 to 0.7). In each step, variance explained values of the selected voxels were bootstrapped 1000 times with replacement. In each iteration, the difference between the median of the two models was computed, as well as the 95% bootstrapped confidence intervals of the difference. Bootstrapping analysis was also used to estimate the beta values of the negative pRF model in the predefined visual and auditory ROIs. This was done to test whether there are differences in the beta values of hearing and deaf participants, and whether these differences are specific to auditory areas. Similar to the variance explained bootstrapping analysis described above, for each participant, voxels were selected at 5 quantile threshold steps, and the beta values were bootstrapped 1000 times with replacement. For each iteration, the median beta value was computed, as well as the 95% bootstrapped confidence intervals.

## Results

This experiment aimed at exploring the organization and spatial properties of cross-modal visual responses in the auditory cortex of congenitally deaf participants, using pRF modelling. During the fMRI session, two deaf and a hearing participant were presented with typical stimuli for retinotopic mapping (**Figure 1)** (Engel et al., 1994, 1997; Sereno et al., 1995). In our modelling strategy we account for both positive and negative responses, thus characterizing positive and negative pRFs.

### Visual representation in the visual cortex

First, we analysed the data using a conventional pRF modelling, in which the selection of the best fitted predictor was restricted to predictors with positive beta values (“positive model”), thus fitting a positive pRF to each voxel. Individual retinotopic maps presented on the inflated cortical surface (**Figure 2**), reveal the known topographic organization of early visual cortex. In both hearing and deaf participants, eccentricity preferences range from fovea to periphery along the anterior-posterior axis (**Figure 2A**). Polar angle maps demonstrate the lateralization of pRFs, as voxels in early visual cortex represent the contralateral hemifield (**Figure 2B**). The reversals in the representation between the vertical and horizontal meridian correspond to the borders between early visual areas (V1, V2, and V3), as delineated based on the HCP MMP atlas (Glasser et al., 2016, **Figure 2B**).

**Figure 2:**
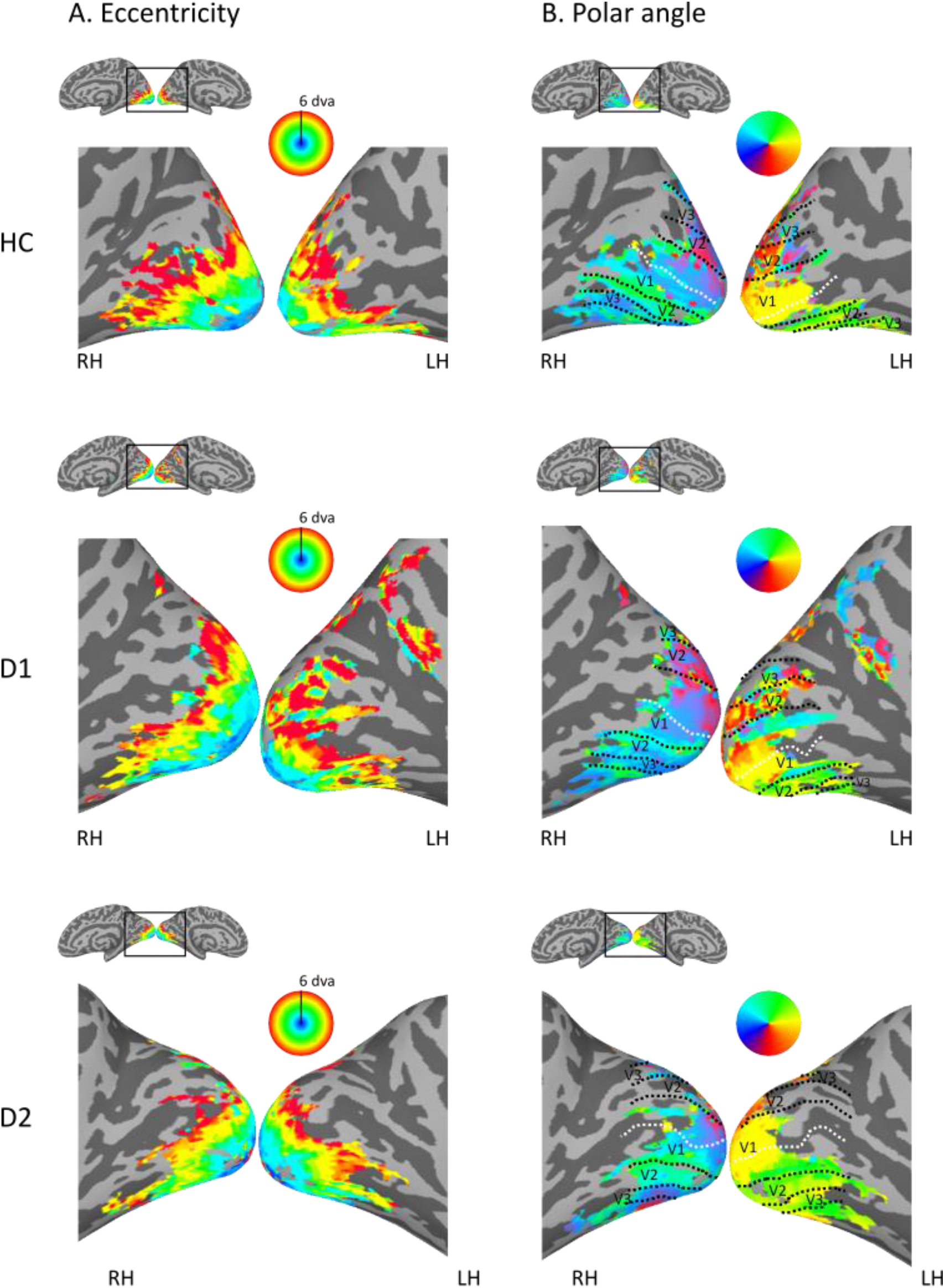
Retinotopic organization in early visual cortex. Eccentricity **(A)** and polar angle **(B)** maps derived from positive pRF modelling are represented along the early visual cortex of hearing control and deaf participants. Threshold was set to R^2^ ≥ 0.20. Dashed lines delineate the calcarine sulcus (white) and borders between V1, V2 and V3 (black lines), that were defined based on the projection of the HCP MMP atlas (Glasser et al., 2016) on each individual cortical surface. HC-hearing control, D1-deaf participant 1, D2-deaf participant 2.

Next, we turned to explore the negative pRF model, and compared the extracted parameters between the negative and positive model. First, we compared the goodness of fit of the two models, as reflected in the variance explained maps. **Figure 3A** shows that both models perform similarly and provide good fits of the data in early visual cortex Mean variance explained values (at a threshold of R^2^ >.0.2) for the positive model and negative models were 0.54 (±0.17) and 0.53 (±1.8) for the hearing participant; 0.62 (±0.18) and 0.59 (±0.20) for participant D1; 0.51 (±0.16) and 0.49 (±1.7) for participant D2. Inspection of the beta values assigned by each model (**Figure 3B**) indicate that in both positive and negative models most responses in the early visual cortex were best fitted by positive predictors. The negative pRF model covers additional areas compared to the positive model, mainly in cuneus and medial parietal areas (see Szinte & Knapen (2020) for a similar result).

**Figure 3:**
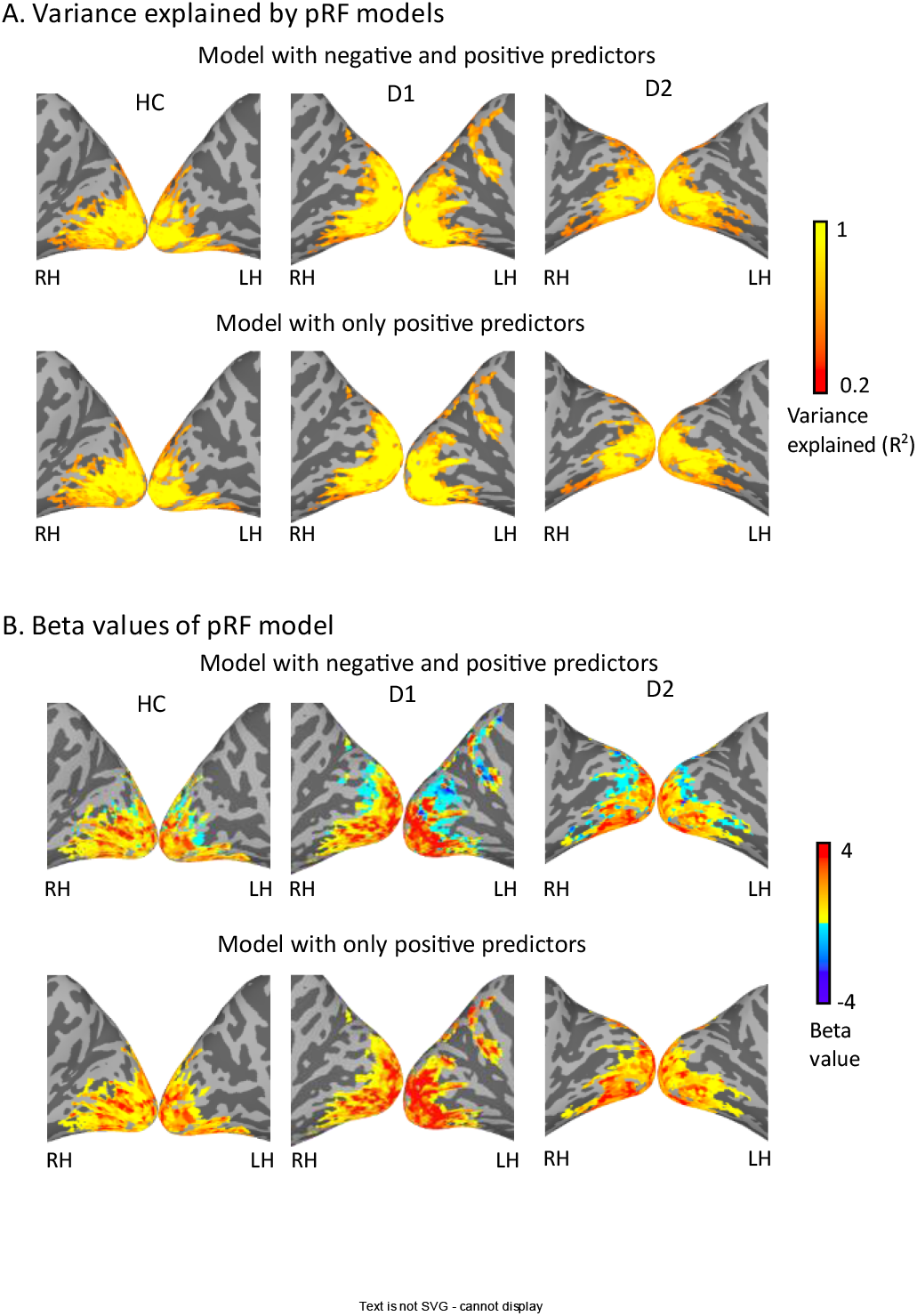
Positive and negative pRF modelling in the early visual cortex. (A) Variance explained by negative (upper row), and positive (lower row) models presented on the medial surface of early visual cortex. **(B)** Beta values of negative and positive models, color coded by their sign. Maps were threshold at R^2^ ≥ 0.2.

To quantitively test whether there is a difference in the performance of the two models, we compared their goodness of fit in the retinotopically organized early visual cortex, as well as in the white matter. The difference in the variance explained (R^2^ values) between the negative and positive pRF models was calculated using bootstrapping quantile analysis at five different thresholds (**Figure 4**; see Methods for ROI definition). The results in the early visual cortex indicate that overall, the two models resulted with similar R^2^ values, except for the lowest thresholds in the deaf participants (**Figure 4**, middle and lower left panels) in which the positive model provided higher R^2^. No differences between the models were found in the white matter ROI.

**Figure 4:**
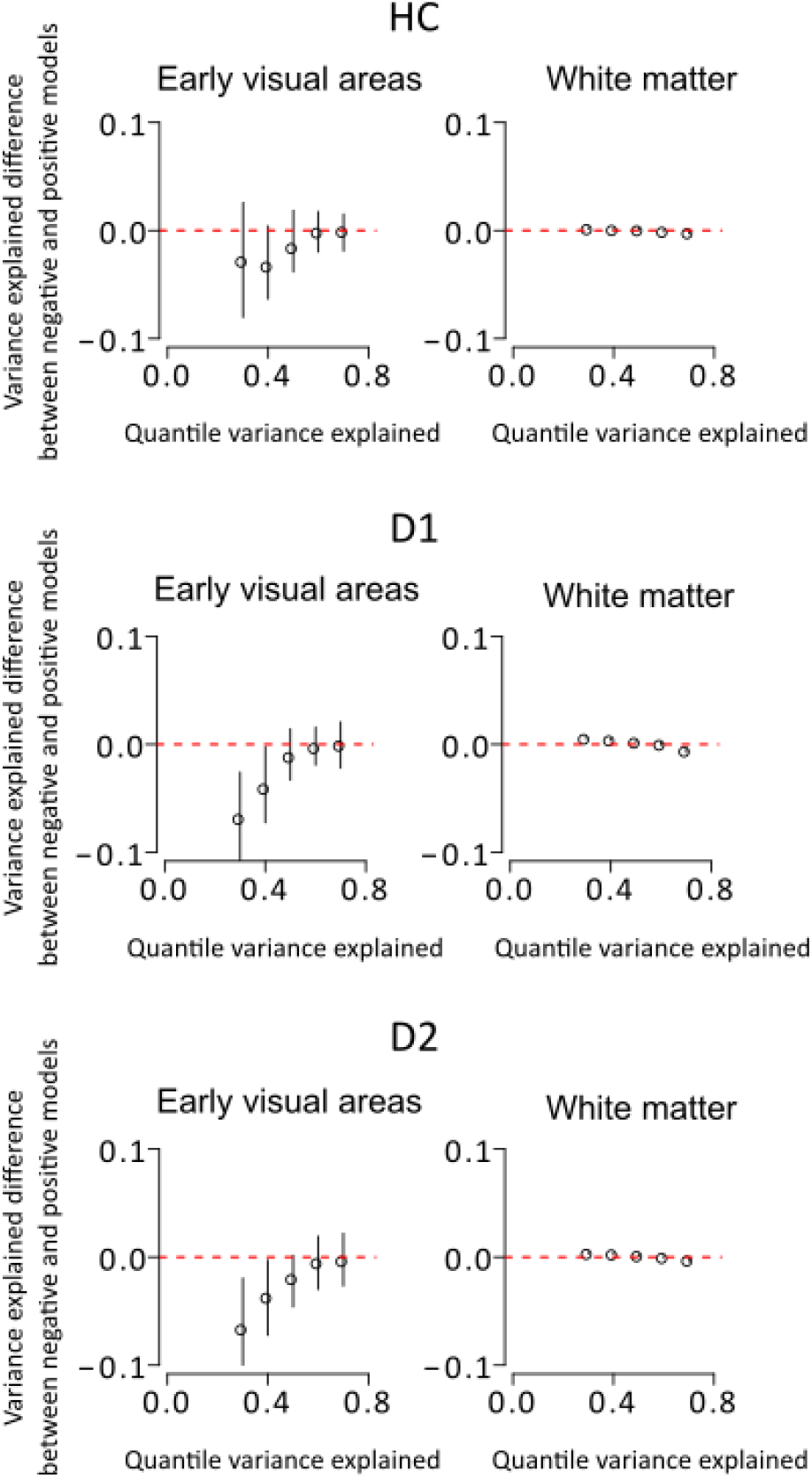
Difference in the goodeness of fit between the negative and positive pRF modelling. The difference (and the 95% confidence intervals) in variance explained values obtained by the two pRF models presented at 5 different quantile thresholds. The two models resulted with similar R^2^ values, except for the lowest thresholds in the deaf participants (D1 & D2) in which the positive model provided higher R^2^.

### Visual representation in the auditory cortex

The positive and negative models provide similar results for deaf and hearing participants when applied to early visual areas. However, differences between models are evident in cortical locations beyond visual areas, and specifically in the auditory cortex. **Figure 5** presents variance explained maps of both models on the lateral view of the brain, at a lower threshold (R^2^ ≥ 0.08), which was set to be higher than the average variance explained by the model in the white matter (Mean white matter R^2^ values were 0.06 ±0.09, 0.08 ±0.13 and 0.06 ±0.1 for participants HC, D1 and D2, respectively). While the hearing participant does not appear to show clear differences between the two models, the maps of the deaf participants reveal more clusters in the frontal and temporal lobe for the model that includes negative beta values.

**Figure 5:**
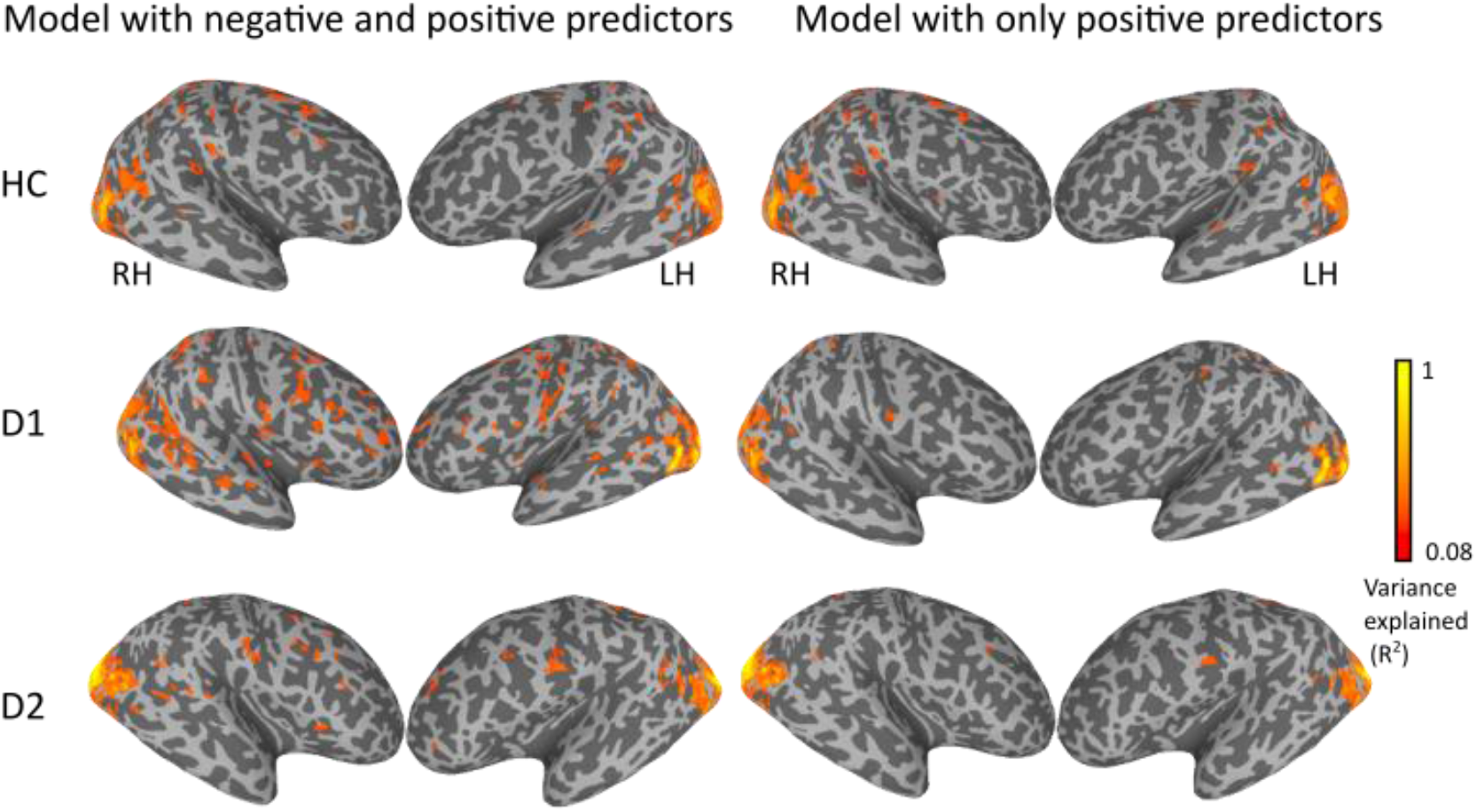
Variance explained by the two models in hearing and deaf participants. Variance explained by the model including positive and negative predictors (left row) and only positive predictors (right row) are presented on the lateral view of the brain. Maps are threshold at R^2^ ≥ 0.08. The maps of the deaf participants (D1 & D2) reveal more clusters in the frontal and temporal lobe for the model that includes negative beta values.

Thus, it seems that potential retinotopic responses in the AC are better captured by the negative pRF model, which led us to an exploratory analysis of these clusters with a focus on this model in the following analysis steps.

To further explore the results obtained by the negative pRF model we plotted the beta values of the best fitted predictor at each cortical location (**Figure 6**). The maps clearly demonstrate that in deaf (but not the hearing) individuals most of the responses beyond visual cortex are characterized by negative betas values, mainly in the frontal and temporal lobes. The difference between the hearing and deaf participants is more evident when plotting the negative and positive values in separate maps (**Figure 6B**). This is particularly visible in participant D1, which shows pertinent clusters in auditory areas including the superior temporal sulcus (STS), superior temporal gyrus (STG), and Heschl’s gyrus (HG), and to a lesser degree in D2. In both deaf participants, additional clusters were also localized to multisensory areas at the temporoparietal junction. **Figure 6C** presents bootstrapping analysis of the estimated beta values of the negative pRF model, which allows for positive as well as negative beta values, in visual and auditory ROIs. The results clearly show that regardless of the chosen threshold, while early visual areas are characterized by positive betas values (in both hearing and deaf participants), the responses in the auditory cortex of deaf participants (and not the hearing participant) are best fitted by negative beta values.

**Figure 6:**
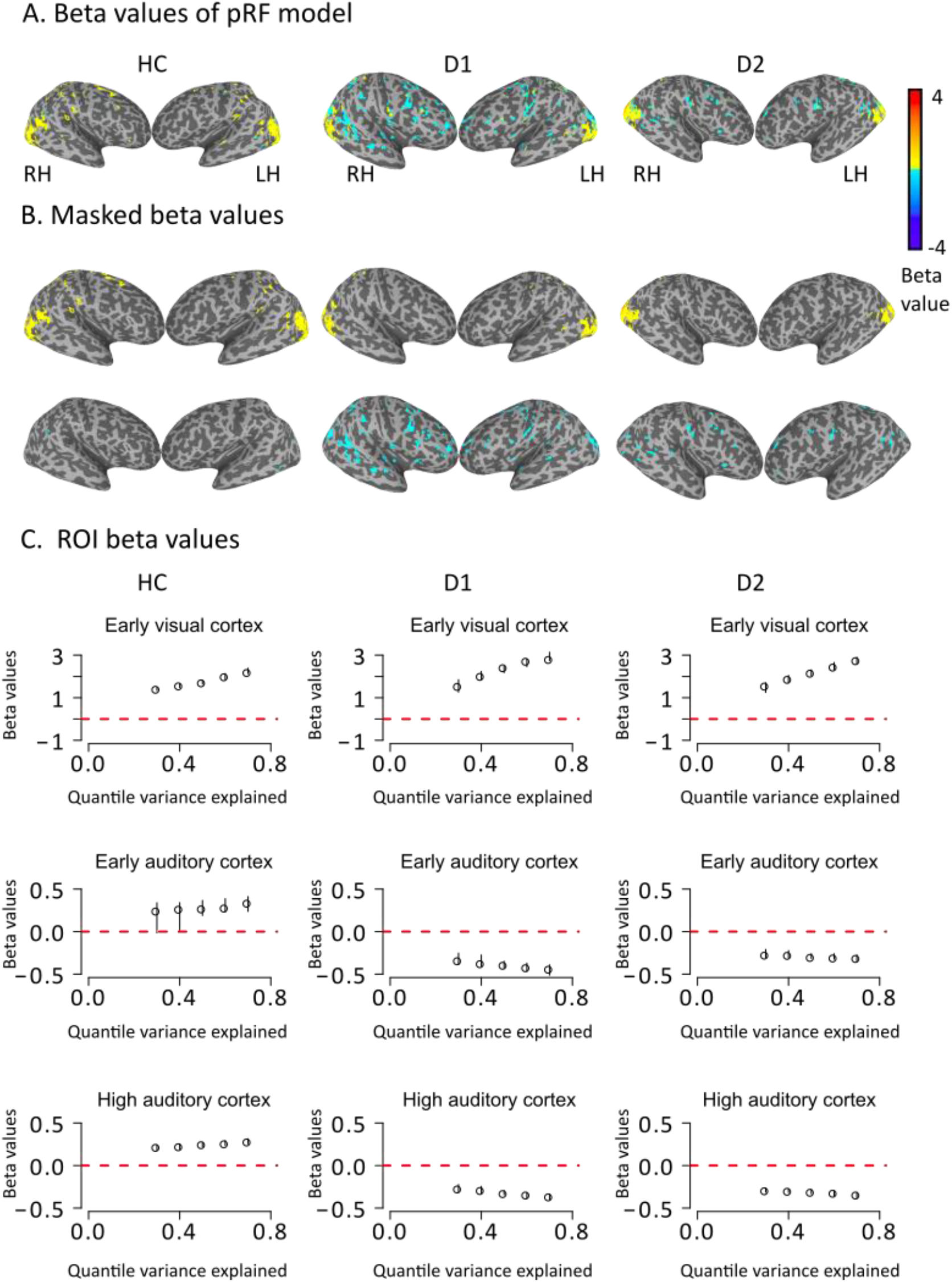
Beta values assigned by the negative model in deaf and hearing participants. Beta values are colour coded according to their sign (warm colours code positive values and cold colours code negative values) and presented as a full map **(A)** or as two separate maps, masked by their sign - warm colours indicating positive beta values, cold colours indicating negative beta values **(B)**. Maps are threshold at R^2^ ≥ 0.08. **(C)** Bootstrapping analysis of the estimated beta values in early visual, early auditory, and higher-order auditory areas. The median beta values (and the 95% confidence intervals) obtained by the negative pRF model are presented at 5 different quantile thresholds.

Finally, we turned to explore which visual features are represented in the auditory areas delineated by the negative model. Because visual responses were seen only in the AC of the deaf participants, and only for negative pRF tuning, we focused on these participants and model in the following analysis. We extracted the parameters representing polar angle, eccentricity, horizontal and vertical location, and pRF size (**Figure 7**) only in voxels with negative beta values. The results reveal that most of the visual-related responses are localized in the right hemisphere, mainly along the posterior and anterior STS and STG. In participant D1 there is also a visible cluster at the level of the Transverse temporal gyrus, corresponding to Heschl’s gyrus (**Figure 7A**). Polar angle maps (**Figure 7A**) show that most of the responses in the right hemisphere represent the contralateral visual field, similar to the known laterality observed in early visual cortex. Eccentricity maps (**Figure 7B**) suggest that the pRFs in AC mostly represent central areas of the visual field, with foveal and near-foveal preferences (values smaller than 2-3 dva). This is further supported by the horizontal and vertical location maps (**Figures 7C** and **7D**, respectively) showing that there is a notable preference for representing the left visual field. In participant D1, right hemisphere, the average horizontal locations values were negative (in the early auditory cortex: mean=-0.63, t(11)=-3.96, p< 0.05, corrected; in the high-order auditory areas: mean=-1.05, t(40)=-3.55, p<0.05, corrected; See **Supp. Table1** for the results of both hemispheres) implying for representation of the contralateral (left) visual hemifield. The average horizontal value in participant D2, right hemisphere, was also negative, but did not reach a significant level (mean= −0.53, t(21)=-2.05, p=0.05). The centre of the pRFs in the right hemisphere were localized near the fixation point in the vertical axis, without a clear preference to the upper or lower visual field. (Mean values for participant D1: −0.11, t(11)= −0.54, p= 0.6 and −0.16, t(40)= −0.88, p=0.39 for the early and high auditory areas. For participant D2: mean=-0.55, t(21)=-1.81, p=0.09). Finally, pRFs size (**Figure 7E**) shows large values, meaning large receptive fields size, in the absence of a consistent gradient. To summarize, the visual responses in the associative and early auditory cortex in participant D1 and D2 show a patchy representation of the contralateral hemifield, with large receptive fields centred near foveal areas.

**Figure 7:**
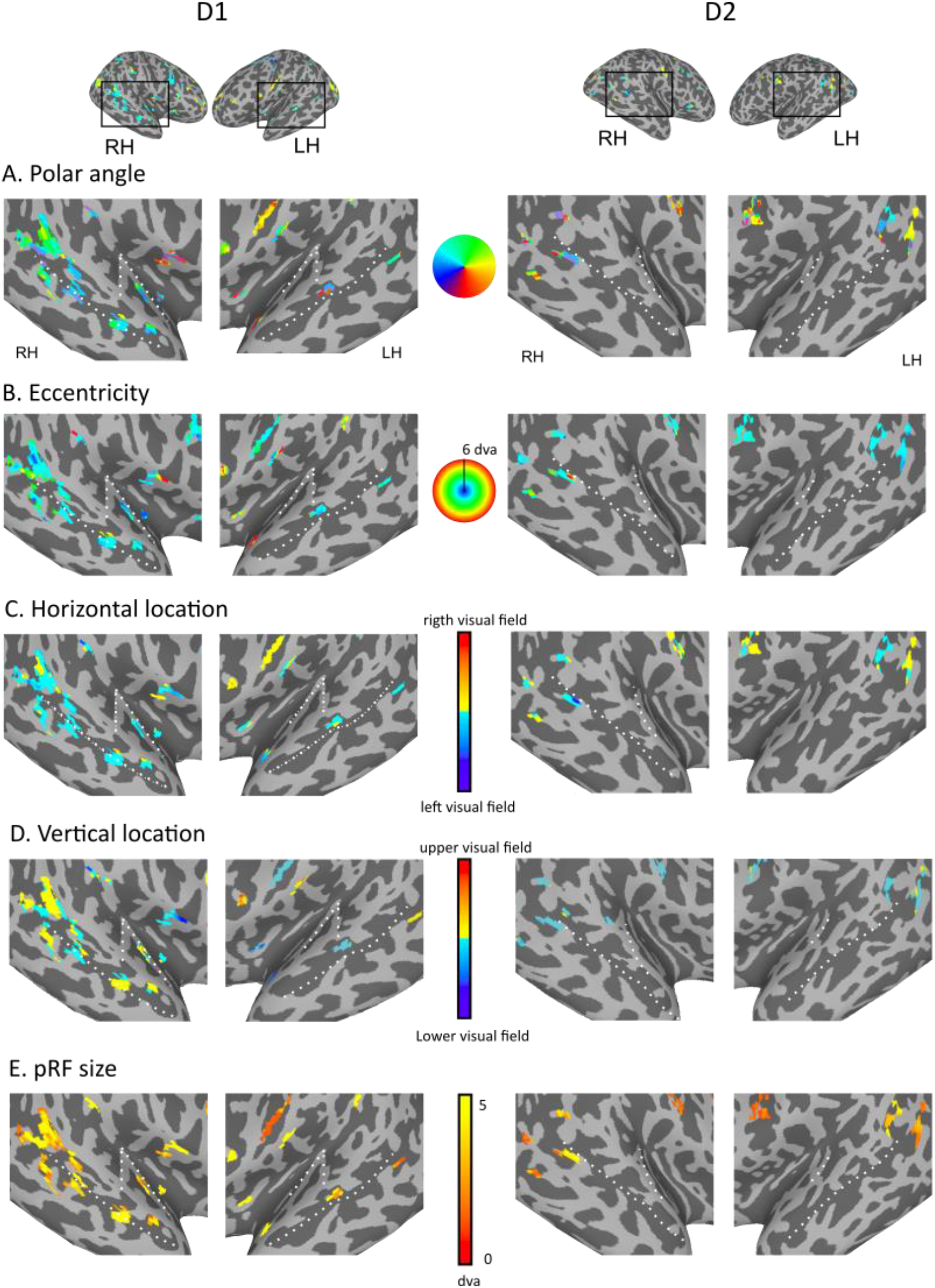
Visual features representations in the AC of the deaf participants. Polar angle **(A)**, eccentricity **(B)**, horizontal location **(C)**, vertical location **(D)** and pRF size **(E)** parameters presented on enlarged view of the temporal lobe of the deaf participants. Maps are threshold at R^2^ ≥ 0.08. Dotted lines delineate the HG and STS.

## Discussion

This study aimed to investigate cross-modal plasticity in congenitally deaf participants by mapping visual responses in auditory areas and characterizing the retinotopic information using population receptive field modelling. The results revealed visual-related responses in the AC of the deaf individuals following passive viewing of simple visual stimulation, but not in the hearing individual. Importantly, our results further show that these visual responses predominantly reflected negative BOLD signals in the auditory cortex of the deaf, suggesting that cross-modal visual information might be represented primarily through deactivation signals.

### Lateralization of cross-modal responses

In the auditory cortex of deaf individuals, the visual responses were mostly localized to the right hemisphere. This lateralization is consistent with current literature on neuroplasticity and known hemispheric asymmetries, indicating that congenital deafness induces structural changes mainly in the right hemisphere (see for example (Hribar et al., 2020; Manno et al., 2021) including alterations in grey matter (Amaral et al., 2016; Tae, 2015), white matter, (Shiell et al., 2016; Shiell & Zatorre, 2017) and subcortical structures (Amaral et al., 2016; Kumar & Mishra, 2018). This structural plasticity was found to correlate with behavioural performance - better performance in visual motion detection tasks was associated with deaf individuals showing increased cortical thickness in the right planum temporale (Shiell et al., 2016) (as well as visual and prefrontal cortices). Moreover, functional MRI studies indicate that right AC is involved in representing visual information in deaf individuals (Almeida et al., 2015; Finney et al., 2001; Seymour et al., 2017; Vachon et al., 2013). The visual-related clusters found in our pRF modelling show lower variance explained values in auditory areas compared to the visual cortex, which is expected for non-visual areas. Importantly, these clusters were found only in the deaf and not the hearing participant, and they appear in regions that have previously been associated with cross-modal plasticity following sensorial deprivation: the HG (in one participant) and STS (Almeida et al., 2015; Bavelier et al., 2001; Finney et al., 2001; Retter et al., 2018; Scott et al., 2014). For example, the neural activation pattern of primary auditory cortex was found to contain spatial information regarding the location of the visual stimuli in deaf (Almeida et al., 2015) while the STS has been associated with neuronal reorganization in deaf, being involved in motion processing and temporal processing for visual and tactile stimuli (Bavelier et al., 2001; Bola et al., 2017; Scurry et al., 2020; Shiell et al., 2015).

### Retinotopic features of cross-modal responses

Our results indicate that the auditory cortex of deaf individual is responsive to visual stimuli. Although these responses do not reveal clear retinotopic gradients fully covering the visual field, they share some retinotopic features: the polar angle maps show that pRFs in the right hemisphere mostly represent information from the contralateral visual field. Functional laterality is further corroborated by the map of horizontal position, supporting this contralateral representation of the visual field in the RH. Regarding eccentricity preference, pRF centres are mostly localized to the foveal or near foveal areas of the visual field, without an apparent bias toward upper or lower visual field representations. pRF sizes are large, thus covering wide areas of the contralateral visual field. Our results are in line with the findings obtained by Almeida et al. (2015) who showed that the quadrant location of a visual stimulus could be decoded from the auditory cortex of the deaf group, and that coding accuracy was higher for the horizontal axis (left versus right hemifields) than for the vertical axis (upper versus lower hemifields). Moreover, classification performance for eccentricity information (central versus peripheral stimuli) was higher than chance level only in the right auditory cortex of deaf group.

Our findings are compatible with single-unit animal studies on cross modal plasticity in auditory areas. For example, the receptive field of visual-responsive neurons in the anterior auditory field (AAF) of deaf cats were large and represented the contralateral visual field (Meredith & Lomber, 2011). Most of the neurons were selective to movement-direction and showed preference for high velocity movement of the visual stimuli. While in some recording sites the spatial organization pattern indicates a progressively shifted location of receptive fields (such as nasal to temporal shifts when the recordings were done oblique with respect to cortical columns), overall, the visual-responsive neurons were sparsely distributed and there was no evidence for a retinotopic map in the sensory deprived areas in the cat (Land et al., 2016; Meredith & Lomber, 2011). A similar pattern was found also for somatosensory cross-modal responses in auditory areas, with large (often bilateral) receptive fields, representing mostly the head and anterior limbs but without a systemic somatotopic gradient (Meredith & Lomber, 2011). Nevertheless, it is important to note that in both humans and animals, cross-modal plasticity processes coexist with a preserved large-scale organization of the deprived sensory cortices, as manifested by functional and anatomical connectivity patterns (Butler & Lomber, 2013; Striem-Amit et al., 2016).

### Negative BOLD and negative pRF

A critical finding in our results is that the visual-related responses in deaf individuals were characterized by negative betas of the pRF model, thus corresponding to below baseline deflection of the BOLD signal in response to the visual stimulus, that is, negative BOLD responses.

Negative BOLD responses in early sensory and motor cortices are usually related to stimuli presented in the ipsilateral visual field (Fracasso et al., 2018, 2021; Smith et al., 2004) as well as ipsilateral hand movement (Hamzei et al., 2002) or tactile stimulation of one side of the body (Kastrup et al., 2008; Tal et al., 2017). Although the physiological basis of NBR remains only partially understood, research has provided some important evidence suggesting that these negative responses reflect a functionally relevant measure of neuronal deactivation, as NBR are correlated with corresponding decreases in cerebral blood flow, oxygen consumption and neuronal activity (Boorman et al., 2010; Mullinger et al., 2014; Shmuel et al., 2002, 2006; Stefanovic et al., 2004). Accumulating evidence have shown that like positive BOLD responses, NBR contain stimulus-specific information, suggesting that these responses contribute to building-up the sensory percept (Bressler et al., 2007; Szinte & Knapen, 2020; Tal et al., 2017; Zeharia et al., 2012). In early sensory cortex, NBR can be found proximal to areas exhibiting positive BOLD responses and were suggested to be mediated by lateral suppression mechanism (Boorman et al., 2010). For example, early visual areas that are tuned to foveal representation respond with positive BOLD signal to foveal visual stimuli and with negative BOLD signal when the stimuli are presented in the periphery, and vice versa (Shmuel et al., 2002, 2006). NBR are also evident in the ipsilateral hemisphere following lateralized visual and somatosensory stimulus or a motor task (Fracasso et al., 2018; Gouws et al., 2014; Hlushchuk & Hari, 2006; Kastrup et al., 2008; Klingner et al., 2010; Tal et al., 2017).

Importantly, NBR have been also reported in cortical areas of the non-corresponding sensory modalities, manifested as negative cross-modal responses, including visual-induced deactivation of the auditory cortex (Laurienti et al., 2002; Mozolic et al., 2008) and negative responses in visual areas following auditory or tactile stimulation (Hairston et al., 2008; Kawashima et al., 1995; Laurienti et al., 2002; Merabet et al., 2007; Sadato et al., 1996; Tal et al., 2016; Weisser et al., 2005). For example, Laurienti et al. (2002) used fMRI in a passive stimulation paradigm with 3 conditions (a visual stimulus alone, an auditory stimulus alone, and a combined visual-auditory stimulus) and concluded that visual stimulation resulted in negative BOLD signals in the AC, namely along the superior and middle temporal gyri with a peak in Brodmann area 22. A similar effect occurred with the auditory condition, as negative BOLD was observed in extrastriate visual areas, but not in response to the combined visuo-auditory condition, which resulted with positive BOLD responses.

Our results provide additional evidence for negative cross-modal responses in the auditory cortex, and further show that these responses are partially tuned to specific parts of the visual field. However, these findings contrast with previous studies of cross-modal plasticity which collectively point to recruitment of the deprived auditory areas for visual or somatosensory tasks (Bola et al., 2017; Karns et al., 2012; Scott et al., 2014; Scurry et al., 2020; Shiell et al., 2015; Vachon et al., 2013). What could be the source for this discrepancy? One explanation could be related to a general bias in the literature for reporting positive modulations of the BOLD signal, rather than BOLD deactivations. For example, the presented maps or statistical measurements in previous cross-modal studies (Benetti et al., 2021; Scott et al., 2014; Shiell et al., 2015; Simon et al., 2020; Vachon et al., 2013) are usually a result of contrast-based analysis between two conditions or groups, and thus do not show the activity pattern compared to baseline. In other cases, cross-modal NBR have been demonstrated (Karns et al., 2012), but these responses were not further characterized. In fact, recent studies suggest that NBR might be much more prevalent although their detection is highly task-dependent (Gouws et al., 2014). Higher task demands lead to a substantial increase in NBR (Gonzalez-Castillo et al., 2015), and the difficulty of the task determines the degree of deactivation (Hairston et al., 2008). Future studies, aiming to explore the prevalence of cross-modal NBR under different behavioural tasks and cognitive load conditions, could provide important insights regarding their role in cross-modal plasticity. Finally, NBR could be detected also in response to passive viewing of stimuli (Mayhew et al., 2013), however, since the amplitude of NBR is substantially lower than the PBR (Gonzalez-Castillo et al., 2015; Harel et al., 2002), it requires a higher signal-to-noise ratio to be detected, which could be achieved by obtaining a large dataset (large groups of participants or the collection of substantially longer fMRI data of single participants) or by using a stronger magnetic field (Jorge et al., 2018). Nevertheless, despite our small-sample size we were able to observe task-dependent NBR at the single subject level, specifically for deaf individuals. Given that all participants were presented with the same stimuli in an equal context, it is reasonable to assume that differences between deaf-hearing arise due to their life-long sensory deprivation and functional and anatomical plasticity.

What could be the functional relevance of negative pRF found in the deaf? As described above, converging evidence suggest that NBR are much more prevalent than commonly reported, suggesting that these responses might be relevant to different aspects of sensory-motor and cognitive processing. In our results, negative pRF were found beyond the auditory cortex in medial areas (cuneus/medial parietal) and prefrontal cortices associated with the DMN, with some similarities to a recent study (Szinte & Knapen, 2020) that explored the role of the DMN in visual processing. DMN areas typically show activation during rest (Raichle et al., 2001) and self-related cognitive tasks such as episodic and semantic memory or mind wandering, and deactivation during attention-demanding and externally oriented tasks (Alves et al., 2019). By allowing the pRF model to capture both positive and negative BOLD responses, Szinte & Knapen (2020) showed that the DMN selectivity deactivates as a function of the position of a visual stimulus. Their study was the first ascribing the term “negative pRFs” to refer to signals that are better predicted by deactivation and are tuned to the spatiotemporal features of to the visual stimuli. Negative pRF were further found in non-human primates, in a study that modelled whole-brain fMRI signal and neurophysiological multi-unit recording from the visual cortex (Klink et al., 2021). BOLD-driven negative pRFs were found in peripheral parts of V1, and in areas that have been associated with the monkey DMN. These findings imply for a sensory-related organization of the DMN that potentially could be selective also to other sensory modalities (Szinte & Knapen, 2020). The dynamic balance between positive and negative activity might possess computational benefits and increase processing efficacy within and across modalities. That is, encoding the dissimilar features might provide an effective approach for perceptual inference (Goncalves & Welchman, 2017). For example, computational modelling approaches have shown that a mixed population of neurons with mismatched tuning for two different sensory cues (such as visual and vestibular movement direction tuning) reduces errors in decoding the possible sources of sensory inputs (Kim et al., 2016).

Although NBR might arise from different neuronal sources under different experimental paradigms and attentional demands (including lateral inhibition in local scale, interhemispheric inhibitory connections, or top-down modulation), these negative modulations occur in response to unattended stimuli, i.e., in areas beyond the corresponding receptive field, or in the unattended sensory modality. Thus, NBR might serve in sharpening the overall neural population tuning and operate as a filter for non-relevant information both within and across sensory modalities. If deactivating/suppressing nonrelevant information contributes to the perceptual integrity and attention to sensory information (Hairston et al., 2008), we speculate that the observed negative pRF in our study might be involved in the enhanced performance of the deaf in some visual tasks mainly in the peripheral visual field (Bavelier et al., 2000; Bosworth & Dobkins, 2002; Neville & Lawson, 1987). We found that in the auditory cortex of deaf pRF centres are located near the fovea, and thus the peak of deactivation arises during foveal and near-foveal stimuli. The suppression of foveal visual field could increase their ability to distinguish between central and peripheral visual representations, in accordance with their enhanced performance in visual peripheral tasks. The fact that the visual-induced negative pRFs in the auditory areas were not accompanied by nearby positive pRFs with peripheral selectivity (as previously found in peripheral V1 areas (Klink et al., 2021; Smith et al., 2004)) suggests that the underlying mechanism is probably less related to local surround-suppression and might involve top-down sensory or attentional modulations. Future studies, with modulated attentional paradigm and higher spatial resolution could further support this hypothesis and uncover the selectivity of cross-modal negative pRF in processing visual information.

## Acknowledgements

This research is supported by an FCT grant PTDC/PSI-GER/30757/2017 and Programa COMPETE Portugal. JA and ZT are supported by funds from the European Research Council (ERC) under the European Union’s Horizon 2020 research and innovation programme Starting Grant number 802553; “ContentMAP” attributted to JA. JS is supported by an FCT grant for doctoral research (UIDP/00730/2020. Lugus 842344). FF is supported by the National Natural Science Foundation of China (31930053). A.F. is supported by a grant from the Biotechnology and Biology research council (BBSRC, grant number: BB/S006605/1) and the Bial Foundation, Bial Foundation Grants Programme Grant ID: A-29315, number: 203/2020, grant edition: G-15516.

## Author contribution

ZT-Software, formal analysis, visualization, writing - original draft, review & editing

JS – Software, formal analysis, visualization, writing - original draft

FF – Investigation, resources

YB – Investigation, resources

JA- Conceptualization, resources, writing - review & editing, supervision

AF- Methodology, software, writing - review & editing, supervision

## Notes

### Competing Interest Statement

The authors have declared no competing interest.

