## Supplementary table 1 for "The neural organization of visual information in the auditory cortex of the congenitally deaf"

**Supplementary table 1: Horizontal and vertical location parameters in auditory ROIs of the deaf participants.**

X and Y Location parameters were extracted from voxels with negative pRF, after applying a threshold of R^2^ > 0.08. Statistics were calculated only for ROIs with at least 10 voxels above the threshold.

Deaf 01, uncorrected p values

|  | **Right hemisphere** | | | **Left hemisphere** | | |
| --- | --- | --- | --- | --- | --- | --- |
|  | **number of voxels** | **X location** | **Y location** | **number of voxels** | **X location** | **Y location** |
| **Early auditory** | 12 | mean = -0.63  t = -3.96  p= 0.0022 | mean = -0.11  t = -0.54  p = 0.5997 | 10 | mean = -0.4  t = -0.56  p = 0.5907 | mean = -1.56  t = -2.05  p = 0.0705 |
| **Higher auditory** | 41 | mean = -1.05  t = -3.55  p = 0.001 | mean = -0.16  t = -0.88  p = 0.3861 | 33 | mean = -0.68  t = -1.4197  p = 0.1654 | mean = -1.37  t = -2.22  p = 0.0334 |

Deaf 02, uncorrected p values

|  | **Right hemisphere** | | | **Left hemisphere** | | |
| --- | --- | --- | --- | --- | --- | --- |
|  | **number of voxels** | **X location** | **Y location** | **number of voxels** | **X location** | **Y location** |
| **Early auditory** | 1 | - | - | 4 | - | - |
| **Higher auditory** | 22 | mean = -0.53  t = -2.05  p = 0.0529 | mean = -0.55  t = -1.81  p= 0.0848 | 6 | - | - |
